# Breakdown of modularity in complex networks

**DOI:** 10.1101/142174

**Authors:** Sergi Valverde

## Abstract

The presence of modular organisation is a common property of a wide range of complex systems, from cellular or brain networks to technological graphs. Modularity allows some degree of segregation between different parts of the network and has been suggested to be a prerequisite for the evolvability of biological systems. In technology, modularity defines a clear division of tasks and it is an explicit design target. However, many natural and artificial systems experience a breakdown in their modular pattern of connections, which has been associated to failures in hub nodes or the activation of global stress responses. In spite of its importance, no general theory of the breakdown of modularity and its implications has been advanced yet. Here we propose a new, simple model of network landscape where it is possible to exhaustively characterise the breakdown of modularity in a well-defined way. We found that evolution cannot reach maximally modular networks under the presence of functional and cost constraints, implying the breakdown of modularity is an adaptive feature.

## I. INTRODUCTION

Complex networks pervade the evolution and organisation of a wide range of systems, from cellular or brain webs to technological graphs. Their structure has important consequences for their stability, resilience and fragility. Some particular properties of these networks are very common, such as the presence of modular organisation (1–4). In modular webs, different subsets of nodes display a higher integration among them than with the rest of the system. This feature allows some degree of segregation between different parts of the network and has been suggested to be a prerequisite for the evolvability of biological webs (5). Within the context of technological evolution, modular structures have been often proposed as a target for engineering design.

Modules are also expected to play a key role in providing a source of specialisation, while their proper interconnection guarantee integration at the system-level scale. Both are needed in order to sustain proper functionality and we need to understand both how modules are generated and how their disconnection leads to functional decay. An illustrative example is provided by brain network topology or the so called *connectome* (6). Connectomics has been a major breakthrough in pushing forward a new approach to brain disease where both brain areas and their connectivity patterns become integrated in a single picture. Under this view, neurological disorders including Alzheimer’s disease or schizophrenia to challenged healthy cognition, such as in sleep or awareness, can be understood in terms of faulty intermodule communication (7–10). This failure can lead to the so called *breakdown of modularity* (BM) first proposed in (11). It involves a transition from high modularity to low modularity. Similar patterns can be found in other areas, but no general theory of this phenomenon and its implications has been advanced yet.

In spite of its importance, BM has received little attention and it is not well-understood. One important reason of this is connected with the difficulties associated to understanding the mapping between structure (genotype) and function (phenotype) in evolved networks. This is specially difficult when dealing with a property as modular structure, and the need for understanding how and when modular networks are expected to evolve and how optimality is tied to modular architecture. For example, it has been suggested that networks evolved under “modularly varying goals” must be modular (17). Specifically, computational experiments showed that optimal networks are non-modular whenever the goal was kept fixed or under randomly varying goals (with no common subgoals). However, Clune and co-workers have shown that modular networks evolve even in the presence of fixed and modular input-output mappings (18). Here modular patterns would be the byproduct of a cost-dependent selection process. More generally, when dealing with evolving networks, an important question is the role played by modular structures in enhancing robust functionalities and how modular structures are associated to evolvable designs. In other words: is the landscape of modularity associated to Boolean functions smooth? Are optimal modular solutions always tied to efficient functions and evolvable architectures?

In order to address these limitations, a simple case study that can be systematically explored would be desirable. Here we propose such a toy model of network landscapes where it is possible to exhaustively characterise BM in a well-defined way. In this context, it is worth noting that BM seems to be a common feature of computational systems (11; 14). Because computation-related networks can be seen as instances of functional Boolean processes performed on well-defined circuits, a minimal case study can help to gain insight into the role played by modular architecture. Specifically, we consider the set of minimal Boolean feed-forward networks implementing the 2^8^ = 256 Boolean functions *f_μ_* with 3 input variables, i. e. the mapping

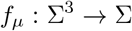

where Σ = {0,1} is the Boolean state space. Such kind of Boolean representation has been widely used in the study of evolved networks and in different contexts, including cellular circuits (12) or pattern-forming genetic circuits (13). A systematic exploration requires necessarily a limitation of the combinatorial space to be analysed. However, relevant computational spaces and specific cases can be observed even in the simplest networks (19). Our analysis suggests that the optimisation of specific inputoutput mappings is not always compatible with highly modular structures and how the BM might be an adaptive feature.

## II. FEED-FORWARD BOOLEAN NETWORKS

The model used here is based on Boolean logic (20), which has been used in the modelling and analysis of the flow of information in natural and artificial systems, such as gene regulatory networks (15; 16). A Boolean function can be represented using a truth table, functional forms and networks and it is worth noting that, despite the Boolean picture is a necessarily simple, neural and genetic networks display nonlinear functional responses that ultimately involve an almost all-or-none behaviour. The interactions between these representations reveal the presence of functional constraints in the organization of complex systems (see below).Our truth tables give the value for the function *f_μ_*(*a*, *b*, *c*) ∈ {0, 1} for each possible combination of the inputs *a*, *b*, and *c*. The function is identified by its designation number

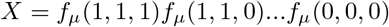

that is, the binary sequence of all function values (see Figure 1A). We can achieve more readable (but ambiguous) expressions using functional forms. The full disjunctive normal form (Figure 1A top) is the sum of elementary products (terms) corresponding to input combinations on which *f_μ_* is true (minterms). For example, the minterm *abc* represents the combination 111, *ab¬c* represents 110 and so on.

**FIG. 1.**
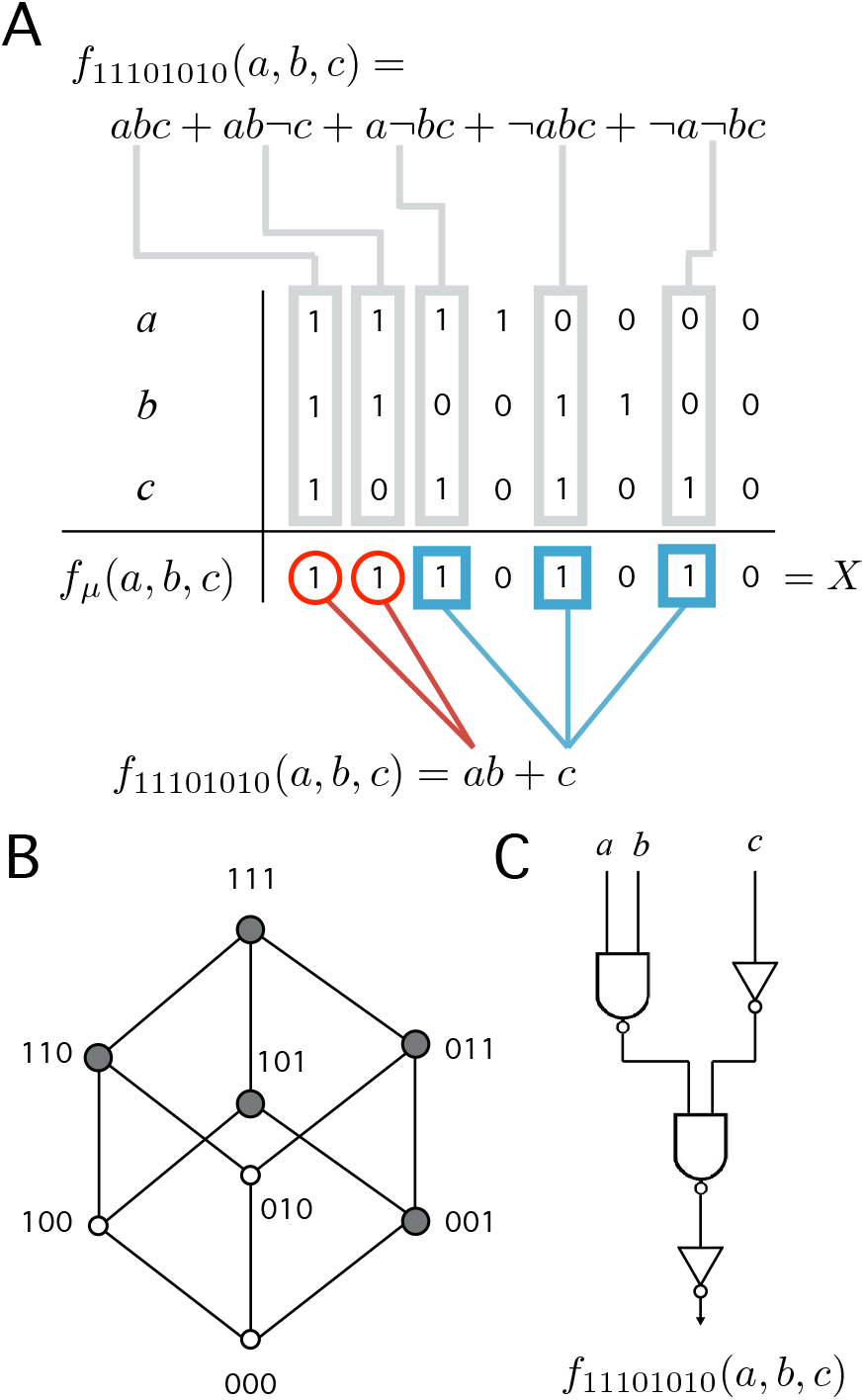
Different representations of a Boolean function. (A) Each function has *ν* = 3 variables named *a*, *b* and *c*. The truth table specifies the value *f_μ_*(*a*, *b*, *c*) ∈ {0,1} for each of the 2^3^ possible combinations of 3 binary inputs (columns). Combinations corresponding to *f_μ_* = 1 are called minterms. A function is identified by its designation number *X* = 11101010 (last row). More readable representations are the full (top) and minimized (bottom) disjunctive normal forms. (B) Hypercube consists of the set of terms (nodes), minterms (gray nodes) and the edges connecting the closest terms in the function. (C) A feed-forward Boolean network (FFBN) is a directed network made up of logic gates and wires that implement the function. The figure shows the FFBN of minimal cost.

A common network representation of a Boolean function is the simple feed-forward Boolean network (FFBN) without no feedback loops. We focus on a subset of the function space *f_μ_* : Σ^3^ → Σ involving all the FFBNs that compute single-output Boolean functions *f_μ_* : {0, 1}^ν^ → {0,1} with *ν* = 3 input variables and one output. The FFBN is a directed graph in which all the nodes carry the labels of negative-AND (NAND) gates while input nodes carry the labels of input variables.

Formally, a directed network *G* = (*V*, *E*) consists of a set of nodes *v_i_* ∈ *V* and a set of edges (*v_i_*, *v_j_*) ∈ *E*. The adjacency matrix *A* = [*A_ij_*] has elements such as *A_ij_* = 1 if there is link (*i, j*) ∈ *E* and *A_ij_* = 0 otherwise. The size of the network is the number of nodes *N* = |*V*| (logic gates) that it contains. The number of links is the sum

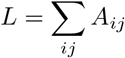

In the following, we will study the undirected version of the FFBN. The network is undirected if for each edge (*v_i_, v_j_*) ∈ *E* there is another edge (*v_j_*, *v_i_*) ∈ *E*. An undirected network has *m* = *L*/2 edges. We also define the degree *k_i_* = Σ_*j*_ *A_i_j* as the number of edges attached to the *v_i_* node. The average degree of the network

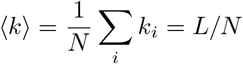

measures the overall connectivity in the system and it is one of the main network parameters.

We will also be interested in measuring the flow of information in the network. The path length is a measure of the distance between nodes in the network. The length l of a path is the number of edges traversed along the path. Let’s define 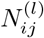 as the number of paths of length *l* that relate any pair of nodes *v_i_* and *v_j_*. Among all the alternative paths, we choose the path of minimal distance (or geodesic path), which defines the shortest path distance *d*(*i*, *j*) or the smallest value of *l* such that 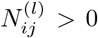. The average path length of a graph (or network diameter) provides a measure of network efficiency and it is defined as follows:

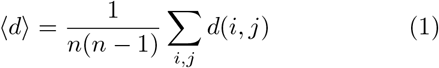

where the normalization term discounts the influence of nodes on themselves.

## III. FUNCTIONAL MODULARITY

The main goal of this paper is to characterise the landscape of Boolean functions associated to our system, and specifically the modularity of neighboring functions and how is the modularity of the circuit implementing each function with its one-bit neighbours. Given a decomposition of the network into a set of subgraphs *C_i_*, the degree of modularity *Q* associated to this partition can be measured as follows (28):

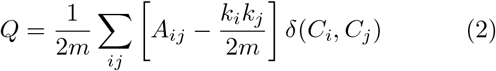

where *k_i_* = Σ_*j*_ *A_ij_* is the number of connections attached to the *i*-th node (or node degree), *C_i_* is the partition the *i* – *th* node belongs to and *δ*(*x*) = 1 is *x* >= 1 and *δ*(*x*) = 0 otherwise. Here, we use the Louvain method for community detection in order to find the optimal partition of the FFBN that maximizes the modularity value (29).

Our hypothesis is that functional requirements constrain structural modularity, that is, evolution cannot reach maximally modular networks under the presence of functional constraints. In this case, we can define a functional (or phenotype) modularity or upper bound for the network (or genotype) modularity (2). However, several genotypes (FFBNs and functional forms) can be found for the same phenotype. When there is representation ambiguity, we often prefer shorter or minimized forms among all the alternatives. For example, a shorter functional form (Figure 1A bottom) divides the support of *f_μ_* into groups (red circles and blue squares) that exploit functional symmetries (30; 31).

Similarly, engineers are often concerned with the problem of obtaining the most economical design for electronic circuits. We define functional modularity *Q*(*X*) as the modularity of the FFBN with minimal cost *L* + *N* or sum of number of logical gates and wires (see Figure 1C). The minimization of Boolean functions is a hard problem (in general) and there is no simple way to obtain the optimal solution (20). Following (21), we perform an exhaustive search to obtain the optimal solution for each of the 256 Boolean functions of 3 variables (see SM for a detailed listing).

## IV. PHENOTYPE NETWORK

In order to uncover the relationship between modularity and functional requirements, we will make use of the concept of phenotypic network. A phenotypic network is a graph whose nodes represent (in our case) Boolean functions and where two functions are connected if they differ in only one minterm of the full disjunctive normal form. In this metagraph, each node maps a function onto its genotype (a FFBN). The notion of phenotypic network is derived from the conceptual framework associated to genotypic and phenotypic spaces proposed by several authors (22–27). The hypercube is also related to the above definition. The hypercube *Q_ν_* is a network of 2^*ν*^ nodes represented by binary sequences 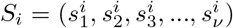 where 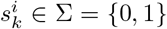. The edges of the hypercube connect nodes whose sequences differ in exactly one bit, i.e., there is an edge (*S_i_*, *S_j_*) ∈ *Q_n_* when *d_H_*(*S_i_*, *S_j_*) = 1. Let’s define the Hamming distance *d_H_*(*S¡, Sj*) between any pair of sequences as follows:

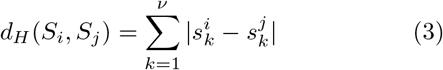

In addition, every Boolean function can be represented as a subgraph of the hypercube (35) (see Figure 1B). Functions in the phenotype network are labelled with its designation number (a binary sequence) and thus, adjacent functions can be formally defined using the Hamming distance between the corresponding designation numbers. The only relevant difference between the standard hypercube and the phenotype network is that the same function can be represented with several nodes (genotypes).

Figure 2 maps the space of all reachable Boolean functions with 3 input variables. Even in this case, which is small and might not seem so relevant, several key classes of functions and circuit designs are involved (see below) either as single networks or as part of larger webs. These maps reveal several interesting features. First, a few functions appear several times in the network because they have more than one minimal FFBNs. For example, the function *f*_10000110_ accepts three different genotypes with the same minimal cost (they are displayed at the bottom left of Figure 2). Second, the network only considers 80 different functions out of the 256 logical functions of 3 variables. Two functions are equivalent and belong to the same class if one can be obtained from the other by a permutation of the input variables. We have discarded from any consideration functions that are equivalent to any class representative (21) (the SM lists the equivalence classes of functions).

**FIG. 2.**
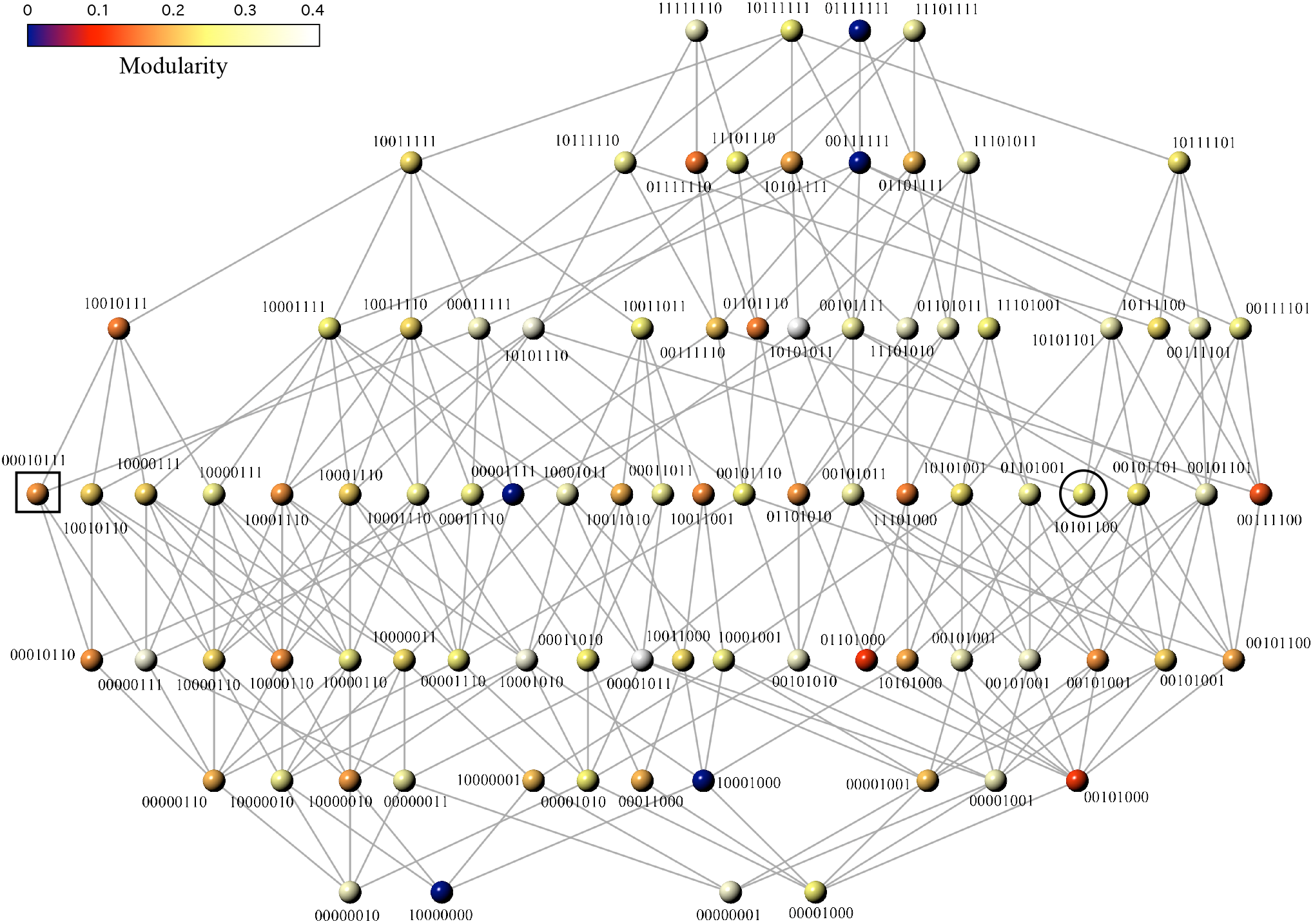
Phenotype network showing all reachable Boolean functions of 3-inputs. Each function *f_μ_* is labelled with its designation number *X* and mapped onto a minimal FFBN. Edges connect pairs of functions *f_μ_* and *f_Y_* with Hamming distance *d_H_*(*X*, *Y*) = 1. Node colour depicts functional modularity, i.e., the modularity value of the minimal FFBN (see text). The black square and circle mark the location of the majority and the multiplexor function, respectively.

## V. ADAPTATION AND THE BREAKDOWN OF MODULARITY

The phenotype network maps the pathways to an eventual BM. Specifically, we can check if it is possible to evolve a less modular target function from any other source function. The distribution of modularity values *P*(*Q*) in this space has a well-defined peak with mean 〈*Q*〉 ≈ 0.2 (see Figure 3A). However, the variance displayed by this distribution suggests the possibility that BM is widespread. Specifically, we can find five functions with minimal modularity (*Q*(*X*) ≈ 0) that can be accessed from different neighbourhoods of the phenotypic space (see yellow region in Figure 3B).

**FIG. 3.**
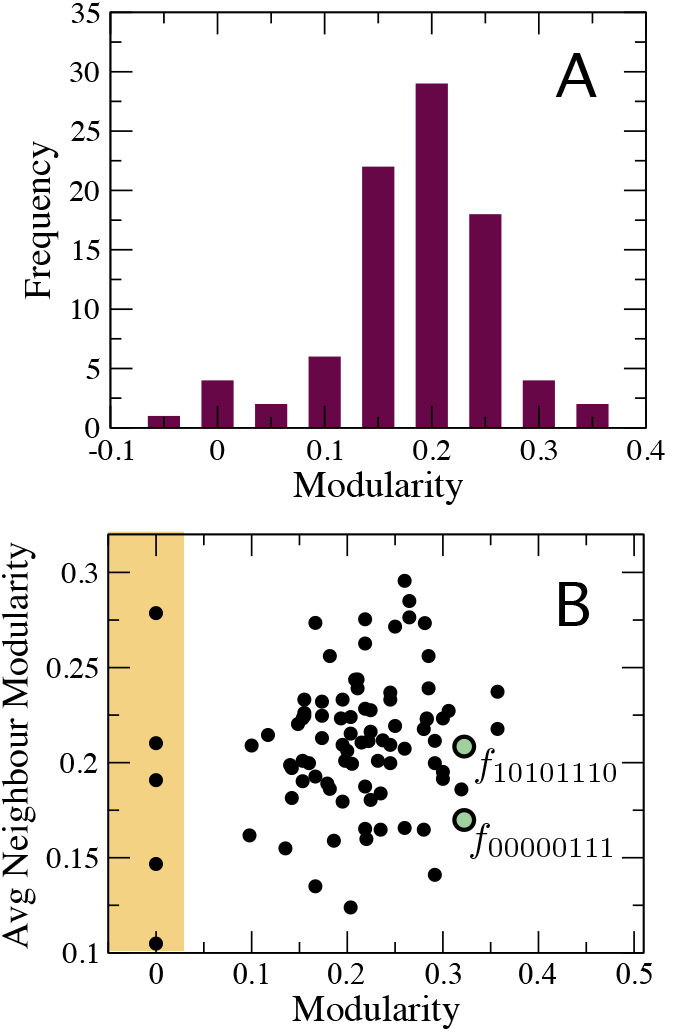
Statistical properties of the phenotype network: (A) distribution of modularity values and (B) correlation between node modularity and the average modularity of its nearest neighbours. The lack of correlation suggests the breakdown of modularity is not a specific property of some systems/environments. Green circles depict the location of two highly modular functions.

There are also highly modular functions surrounded by modular neighbours. To illustrate this behaviour, we have chosen two important functions, namely the multiplexer (see Figure 4A) and the majority function (see Figure 4B). Both circuits have special relevance in both electronic designs and in synthetic biology (? ) Each of these functions can be accessed in a few mutation steps from functions with higher modularity, i.e., *Q*(10101110) ≈ 0.3 and *Q*(00000111) ≈ 0.3, respectively. At least in these two cases, the evolution of useful functions is coupled to a reduction in modularity. These examples suggest how adaptation might lead to a BM. A decrease of functional modularity takes place when the evolutionary goal is non-separable, i.e., the computation of the output requires the interaction of several inputs. For example, the majority function is a global computation that combines all input variables to obtain the output. Similarly, consciously effortful tasks (like working memory) are expected to break the modularity of neural systems (38).

**FIG. 4.**
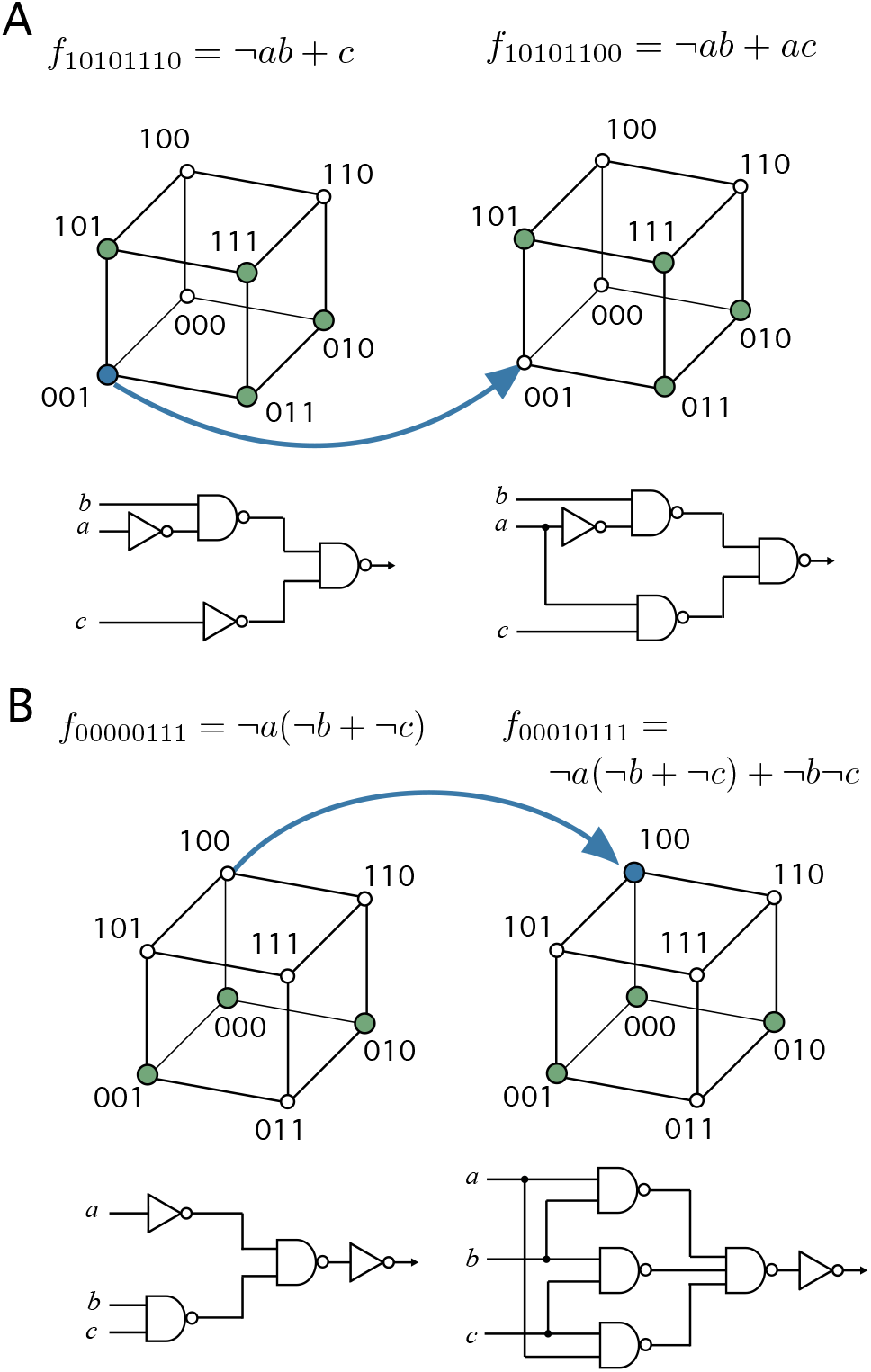
Evolving specific functions might involve a breakdown of modularity: (A) the multiplexer function has functional modularity *Q*(10101100) < *Q*(10101110) and (B) the majority function has *Q*(00010111) < *Q*(000000111). These target functions can be evolved from more modular ancestors, that is, functions with specific groups of inputs affecting different groups of outputs. Blue arrows represent edges in the phenotype network or the specific changes transforming the source function into the target function. Each function is represented with their minimal normal disjunctive form (top), hypercube (middle), and minimal FFBN (bottom).

How can we relate functional characteristics with the breaking of modularity? Figure 3B shows the absence of a clear correlation between the modularity of specific Boolean functions and the average modularity of its nearest neighbours in the phenotype network. Additional network analyses might be helpful to understand this pattern. In a previous study, we have suggested that the BM is related to the small-world behavior of complex networks (11). Software projects have a natural tendency to become disordered structures (41; 48). This degradation is caused by widespread software changes and indirect dependencies between unrelated pieces of code.

These changes in modular organisation have important implications for both engineering and evolved circuit designs. In this context, it was early suggested that software design is an instance of a multi-objective optimization process (36). When designing softare, there is a tradeoff between efficient communication and separation of functional tasks (i.e., modularization). Indeed, small-sized software architectures are trees (as one should expect from optimization leading to hierarchical structures) but clustering emerges at larger sizes. As the number of components increases, conflicting constraints arise between different components that would prevent the reaching of an optimal solution. A need to exchange information between distant parts of a system can lead to a modularity reduction.

The minimization of FFBNs is a similar problem. We can check there is a positive correlation between the average path length and the modularity of minimal FFBNs (see Figure 5A). Modularity leads to an enlargement of the network diameter because there are more pairs of distant nodes accross modules than within modules, that is, we can associate the breakdown of modularity to a reduction of the average path length (1) in the network. For example, the addition of a link between the two branches of the multiplexer (see Figure 4A) leads to a sharp decrease of its average path length. Interestingly, highly modular functions are those having the minimal number of dependencies between the variables (and thus low average degree). Highly modular networks have a tree-like architecture (37) while the presence of functional constraints creates a larger density of connections and a more regular (or lattice-like) topology (see Figure 5). That is, the relationship between network efficiency and modularity reflects the trade-off between function and robustness and evolvability.

**FIG. 5.**
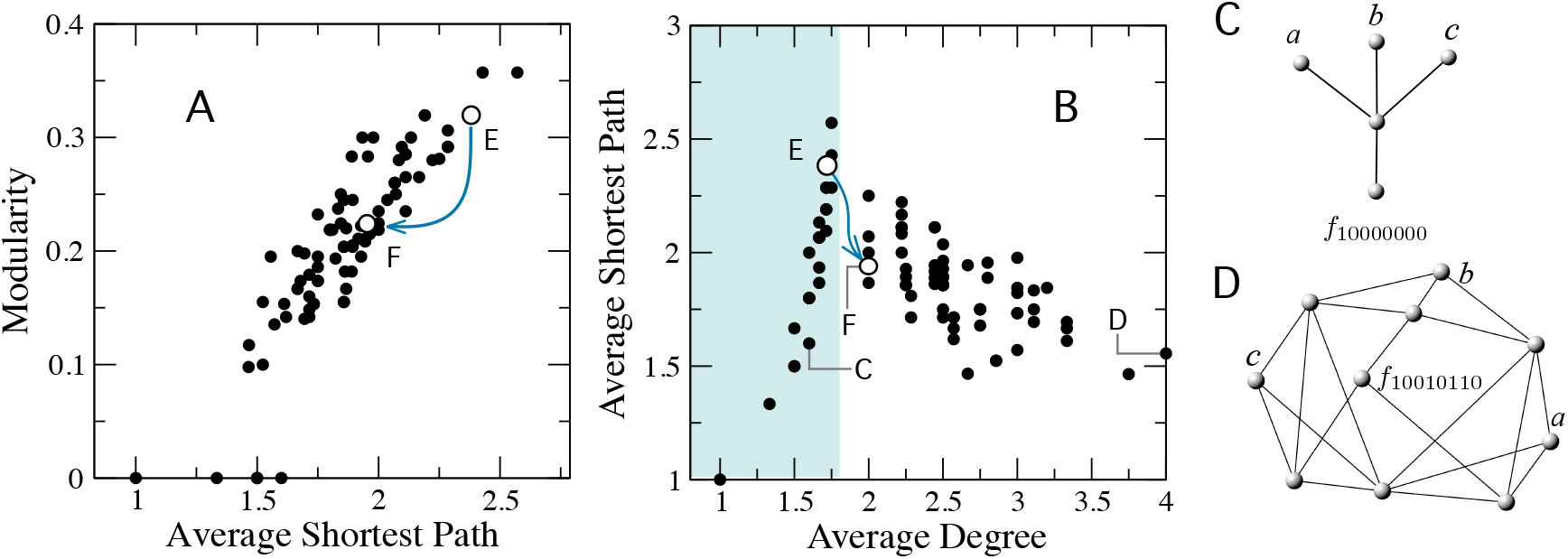
Balance between network diameter and modularity in Boolean networks. (A) Correlation between network diameter and modularity indicates that network diameter has to be sacrified to obtain modular networks.(B) The average degree 〈*k*〉 separates the functional space in two main classes according to their minimal FFBNs showing tree-like (〈*k*〉 ≤ *k*_*_ ≈ 1.75, cyan area) or regular topology 〈*k*〉 > *k*_*_). The minimal FFBNs for the functions *f*_10000000_ (C) and *f*_10010110_ (D) and the location of the multiplexer (F) and the ancestor function (E) are also shown. Blue line in (A) and (B) correspond to the edge (*f*_10101100_, *f*_10101110_) in the phenotype network.

## VI. DISCUSSION

The interplay between fitness and system-level properties such as modularity has been investigated in natural and artificial designs. Simon proposed that nearly decomposable systems composed by independent modules allow faster adaptation to highly fluctuating environments (39; 40). A modular architecture allows independent changes in different parts of the system without affecting the whole. Well-adapted modules are conserved and provide a robust infrastructure for future adaptation. This poses a puzzle because modular architectures cannot be always maintained or reached in artificial evolution. For example, evolutionary algorithms often yield designs that are not decomposable and it is difficult to understand the way these systems work. In software engineering, even if a modular design is provided as initial solution, development rapidly moves to entangled and monolithic solutions (41).

Here we have proposed that the breakdown of modularity takes place because there are changes in the fitness function, e.g., a shift from a well-known environment to a less predictable environment. There are functional constraints to network modularity. For example, tasks involving non-separable input-output mappings like learning a color naming task with interference, evolving a robust metabolic network in a highly fluctuating environment (47) or developing software under constantly changing requirements do not seem to evolve modular networks. Everything is a novelty for a network exposed to a highly fluctuating environment and thus, it makes little sense to maintain costly memories for reusing past information. In a highly fluctuating environment, the only requirement for survival is to issue fast responses and quick adaptations. The lack of memory imposes a strong constraint on the complexity of evolved structures.

The reduction of modularity has to be contrasted with existing theories for the emergence of modularity (51). For example, the breakdown to modularity does not require a change from a modularly changing environment (17) to another static environment. Introducing an additional, temporal dimension in the fitness function is likely to internally decouple the system (and thus creating the possibility of increasing modularity). Moreover, looking at a few case studies does not enable a full understanding of how and when modular networks are expected to evolve. Instead, we have proposed a simple model of network landscape where it is possible to exhaustively characterise the breakdown of modularity in a well-defined way and studied the influence of functional and cost constraints.

The study of biological networks requires the examination of the interactions between modularity, network diameter and function. The phenotype space can be classified in two types of functions depending if the minimal FFBNs (genotypes) is a sparse or a dense network. Sparse genotypes have treelike topologies and their computation involves minimal input interaction. On the other hand, the modularity of dense networks drops with the increasing number of distant interactions. Some of the sparse networks are also maximally modular in our system, contradicting the intuition that modularity depends on densely intra-connected communities. It can be shown that tree modularity is significant even when they are made of sparse modules (37). Nodes in sparse trees acting as bottlenecks are sufficient to achieve high modularity values. The above suggests that we have to extend our definitions of modularity to take into account different measures of internal network connectivity.

Finally, our results might be useful to understand the limits of the hypothesis put forward by Simon. The analysis of the phenotype network reveals how the breakdown of modularity is more likely to take place from regions of the landscape populated by highly modular functions. Although neutral models suggest that tinkering increases the possibilities to discover such modular designs (49; 50), the structure of the phenotype network suggests that it is not always possible to avoid the breaking of modularity. Sometimes modularity is not so beneficial. The evolution of novelties (52) requires the integration of many different sources of information in order to obtain coherent system changes.

## Acknowledgments

I thank Ricard Sole for a careful review of this paper and useful comments and discussions. This work was supported by Botin Foundation by Banco Santander through its Santander Universities Global Division, FIS2016-77447-R from Spain Ministerio de Economía, Industria y Competitividad, AEI/MINEICO/FEDER and UE.

